# Cellular dsRNA interactome reveals the regulatory map of exogenous RNA sensing

**DOI:** 10.1101/2024.06.14.599134

**Authors:** JinA Lim, Namseok Lee, Seonmin Ju, Jeesoo Kim, Subin Mun, Moonhyeon Jeon, Yong-ki Lee, Seok-Hoon Lee, Jayoung Ku, Sujin Kim, Sangsu Bae, Jong-Seo Kim, Yoosik Kim

**Author notes:** These authors contributed equally (JinA Lim, Namseok Lee, Seonmin Ju). Correspondence (J.-S.K.), (Y.K.).

## Abstract

RNA-binding proteins (RBPs) provide a critical post-transcriptional regulatory layer in determining RNA fate. Currently, UV crosslinking followed by oligo-dT pull-down is the golden standard in identifying the RBP repertoire, but such method is ineffective in capturing RBPs that recognize double-stranded RNAs (dsRNAs). Here, we utilize anti-dsRNA antibody immunoprecipitation followed by quantitative mass spectrometry to comprehensively identify RBPs bound to cellular dsRNAs without any external stimulus. Notably, our dsRNA interactome contains proteins involved in sensing N^6^-methyladenosine RNAs and stress granule components. We further perform targeted CRISPR-Cas9 knockout functional screening and discover proteins that can regulate the interferon (IFN) response during exogenous RNA sensing. Interestingly, depleting dsRBPs mostly attenuate the IFN response, and they act as antiviral factors during human β-coronavirus HCoV-OC43 infection. Our dsRNA interactome capture provides an unbiased and comprehensive characterization of putative dsRBPs and will facilitate our understanding of dsRNA sensing in both physiological and pathological contexts.

## INTRODUCTION

Double-stranded RNA (dsRNA) is a common pathogen-associated molecular pattern (PAMP) that is generated as a by-product during RNA virus replication.^1,2^ To counteract viral invasion, cells have developed a collection of pattern recognition receptors (PRRs) that can recognize and bind to dsRNAs to elicit an innate immune response.^3^ Examples of well-known PRRs in mammal cells include melanoma differentiation-associated protein 5 (MDA5), retinoic acid-inducible gene I (RIG-I), protein kinase R (PKR), and toll-like receptor 3 (TLR3).^4,5^ Upon activation, these dsRNA sensors initiate several cellular responses, such as type I interferon (IFN) production, dsRNA degradation, and translational repression, collectively suppressing viral replication.

Recently, several reports suggested that the human genome includes a large portion of repeat elements whose RNAs can also adopt a double-stranded secondary structure.^6–9^ Some abundant dsRNA-generating repeat elements include short interspersed nuclear elements (SINEs), long interspersed nuclear elements (LINEs), and endogenous retroviruses (ERVs). In addition, mitochondria also generate a substantial amount of cellular dsRNAs via bidirectional transcription of the mitochondrial circular genome.^10^ More importantly, these endogenous dsRNAs can interact and activate PRRs just like their viral counterparts, which can lead to fatal consequences.^11–13^ For example, mitochondrial dsRNAs are elevated in synovial fluids of osteoarthritis patients, tear and saliva of autoimmune Sjögren’s disease patients, and blood of Huntington’s disease patients.^14–16^ Moreover, dysregulation and misrecognition of SINE dsRNAs are associated with the development of Aicardi-Goutières syndrome and age-related macular degeneration.^17–19^ Thus, understanding the regulation of cellular dsRNAs holds a key in elucidating the pathogenesis of inflammatory and degenerative diseases.

Aberrant immune activation to cellular self-dsRNAs occurs as these RNAs adopt an A-form helical structure with a deep and narrow major groove that prevents the access of individual nucleotides to RBPs. Consequently, innate immune dsRNA sensors typically recognize the length and structural features, such as 5’ triphosphate, to distinguish self from non-self RNAs.^3,20,21^ More importantly, such sequence-independent interaction mechanism is shared by other dsRNA-binding proteins (dsRBPs), which result in most of them binding to a similar pool of dsRNAs, such as SINEs and ERVs.^22–27^ As a result, multiple dsRBPs will compete for access to their common target dsRNAs,^28^ and they work together to regulate the downstream immune response to the increased levels of cellular dsRNAs. We have recently shown that Staufen1 (STAU1) binds to ERV and SINE dsRNAs and stabilizes them to enhance the downstream IFN response.^25^ Moreover, adenosine deaminase acting on RNA 1 (ADAR1) and DExH-Box Helicase 9 work together to prevent the recognition of self-dsRNAs by innate immune sensors in breast cancer cells.^29,30^

Despite the pathological relevance and therapeutic potential of cellular dsRNAs, the repertoire of dsRBPs remains largely elusive. The current golden standard of systemic identification of RBPs relies on ultraviolet (UV) crosslinking and pull-down with oligo-dT beads.^31–33^ However, this method cannot be extended to the dsRNA interactome due to low UV crosslinking efficiency between the dsRNA and its binding protein.^34,35^ In addition, dsRNAs are mostly generated from repeat elements that lack poly(A) tails, making oligo-dT pull-down inadequate to enrich these RNAs. A recent study utilized biotin-conjugated polyinosinic-cytidylic acid (poly(I:C)) to discover dsRBPs, but it is unclear whether these proteins can bind to cellular dsRNAs and regulate dsRNA sensors in uninfected states.^36^ Considering that dsRNAs are abundantly expressed in infected cells, utilizing viral RNA as a bait to retrieve viral RNA interactome may provide insights into potential dsRBPs.^37–39^ However, this approach is still limited because viruses develop various mechanisms to shield their dsRNAs from host proteins in order to evade innate immune response.^40–42^ Moreover, such method retrieves the virus-specific list of interactors, and it is unclear whether the interactome binds to single-stranded RNAs (ssRNAs) or dsRNAs.

In this study, we report an unbiased analysis of the endogenous dsRNA interactome in proliferating human embryonic kidney 293T (HEK293T) cells with the goal of analyzing its significance in the modulation of IFN response during exogenous RNA sensing. To this end, we employed an original approach that involved co-immunoprecipitation (co-IP) with an anti-dsRNA K1 antibody^43^ followed by quantitative mass spectrometry. To reduce false positives, we performed quantitative mass spectrometry analysis with two additional conditions: 1) treatment with RNase T1 to degrade ssRNAs, and 2) pull-down with synthetic dsRNA, poly(I:C). We then further assessed the regulatory potential of these putative dsRBPs by implementing targeted CRISPR-Cas9 knockout (KO) screening upon introduction of exogenous dsRNA. Subsequent exploration of viral infection within the dsRNA interactome unveiled previously unrecognized dsRBPs that mitigate viral replication. Collectively, our methodology presents valuable resources in studying dsRNA regulation in both physiological and pathological states.

## RESULTS

### In vitro capture of putative dsRBPs in HEK293T

Previous studies by our and other groups revealed that SINE RNAs occupy the most abundant class of cellular dsRNAs.^44–46^ Considering that these SINEs are about 300 nucleotides long while a typical dsRNA binding protein recognizes RNAs with 10∼20 bp in length,^47–49^ we hypothesized that some of the double-stranded regions must be exposed even after the RNA is bound to dsRBPs. To capture these dsRBPs systematically, we thus utilized a K1 antibody that recognizes dsRNAs longer than 40 bp. Our proposed strategy is presented in Figure 1A where we performed IP using magnetic protein A bead coated with K1 antibody and analyzed the co-IPed proteins via quantitative mass spectrometry.

**Figure 1.**
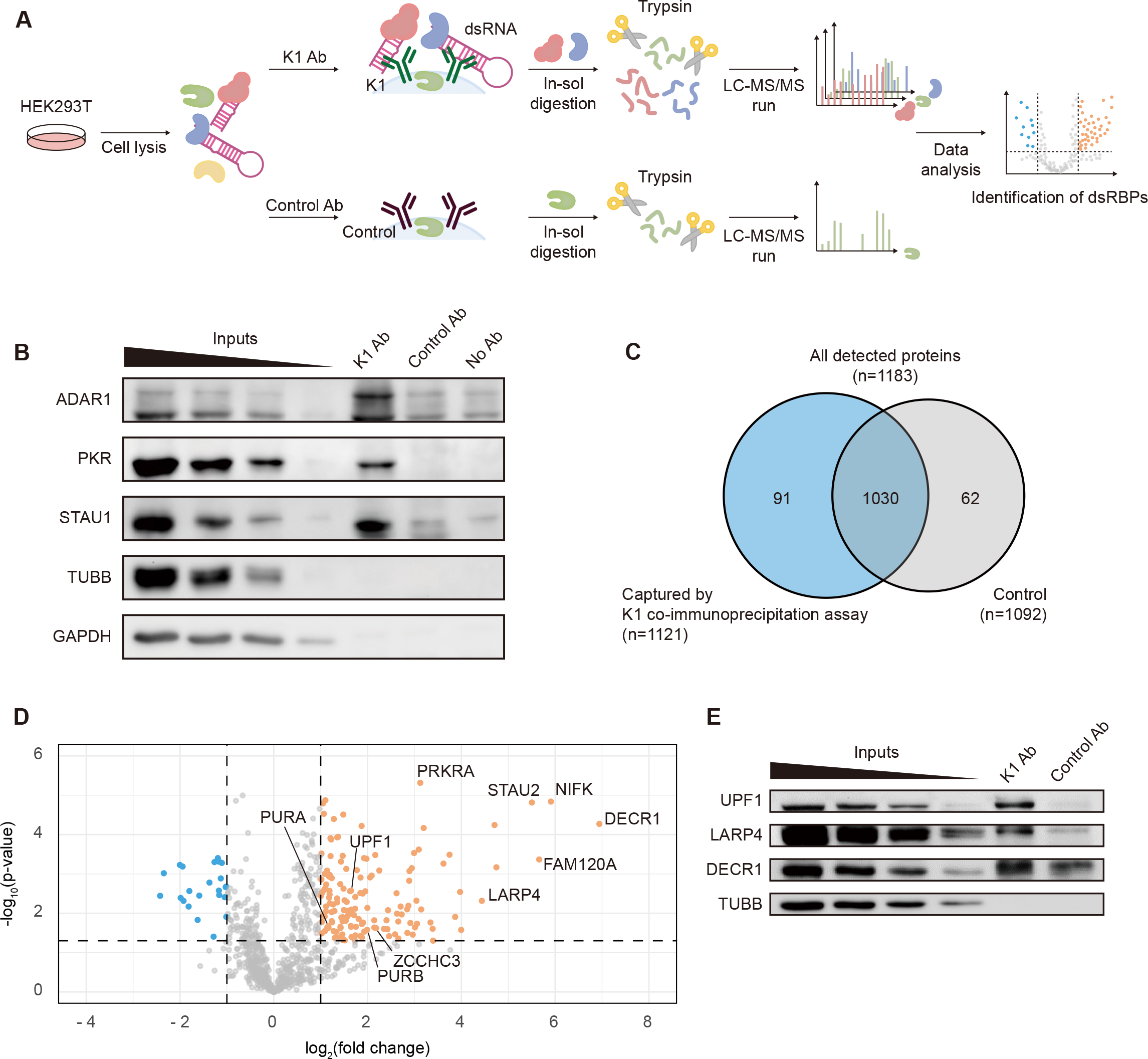
In vitro capture of dsRNA interacting proteins with K1 antibody. (A) Schematics of the K1 IP followed by LC-MS/MS to identify proteins bound to cellular dsRNAs. (B) Western blotting analysis comparing K1 and control antibody captured protein samples. (C) Summary of the number of proteins identified in K1 or control antibody capture. (D) Volcano plot showing proteins that satisfy the filtering criteria: i) identified in all four K1 captured replicates, significantly enriched (orange) proteins with ii) log_2_(fold change) ≥ 1, iii) p-value < 0.05. Log_2_(fold change) was calculated as the difference between averaged log_2_(LFQ intensity) of 4 replicates in each experimental group. Statistical significances were calculated using two-sided Student’s t-tests. (E) Western blot for selected proteins identified through K1 capture that were not previously reported as dsRBPs.

We began by testing the feasibility of our approach using well-known dsRBPs as positive control readouts. We performed an IP experiment with the K1 antibody and analyzed the protein eluate via western blotting. We found that ADAR1, PKR, and STAU1 are strongly enriched in K1 co-IPed eluate, whereas two abundantly expressed non-RBP controls, β-tubulin (TUBB) and glyceraldehyde 3-phosphate dehydrogenase (GAPDH), were undetected (Figure 1B). Moreover, the enrichment of the three tested dsRBPs was specific to K1 co-IPed eluate as they were weakly detected in animal-matched control and no antibody control eluates (Figure 1B).

Encouraged by these results, we performed a large-scale K1 co-IP experiment for liquid chromatography-tandem mass spectrometry (LC-MS/MS) analysis. We identified 1,183 proteins, of which 1,121 belonged to the K1 captured sample and 1,092 to the control sample with animal-matched IgG, with 1,030 (87%) overlapping proteins (Figure 1C). Statistical analysis showed good correlations among biological replicates and distinct characteristics between K1 and control groups in the PCA plot, adding to the reliability of our experimental results (Figures S1A and S1B). When we computed the enrichment for individual proteins, a substantial number of proteins showed positive fold change, indicating that these proteins are likely to be specifically enriched in K1 co-IPed eluates (Figure 1D).

Among the K1 co-IPed proteins detected in all four biological replicates, we selected 148 proteins (p-values of less than 0.05 and a fold change over two compared to control groups) for further analysis (Figure 1D, marked in orange). Interestingly, the protein activator of interferon-induced protein kinase EIF2AK2 (PRKRA), which contains two canonical dsRNA binding domains (dsRBDs), and STAU2, a paralog of STAU1, are strongly enriched, further confirming the reliability of our results. Of note, most of these proteins were not identified by the previous oligo-dT pull-down method, even when formaldehyde was used as a crosslinking reagent.^33,50,51^ In addition, many RNA-associated proteins were specifically pulled down with the K1 antibody. For example, up-frameshift suppressor 1 homolog (UPF1) and La-related protein 4 (LARP4) that can bind to structured elements in the 3’ UTRs were found. An unexpected finding was 2, 4-dienoyl-CoA reductase 1 (DECR1), a mitochondrial metabolic enzyme involved in β-oxidation, as one of the most significantly enriched proteins. As none of these proteins were previously associated with dsRNA-binding, we confirmed whether these proteins are strongly and specifically pulled down by the K1 antibody via K1 IP followed by western blotting. Consistent with the LC-MS/MS results, our validation data clearly showed that the examined proteins are all enriched in the K1 co-IP eluate (Figure 1E).

### Characterization of K1 captured proteins

To analyze the biological characteristics of the proteins captured by the K1 antibody, we first conducted a gene ontology (GO) analysis on molecular function. Consulting GO annotations of 148 proteins revealed that 14 proteins were annotated with ‘dsRNA binding’ (Figure 2A). A significant portion of the remaining proteins (115 out of 134) was labeled with ‘RNA binding’, and 16 proteins were not previously associated with RNAs. Of note, many of these 148 proteins are related to ribosomes, which is likely due to the K1 antibody recognizing ribosomal RNAs (rRNAs). Indeed, while optimizing the K1 capture method, we noticed that many of the K1 captured proteins were ribosomal proteins. This is not surprising, considering rRNAs make up 80% of the RNA present within cells, which would result in some rRNAs inevitably being pulled with dsRNAs, potentially leading to the inclusion of ribosomal proteins in the results. Therefore, we removed small and large ribosomal subunit proteins as well as mitochondrial ribosomal proteins (35 proteins) from our list for further analysis (Figure 2B). We denoted the remaining 113 proteins as the “K1 interactome” (Table S1).

**Figure 2.**
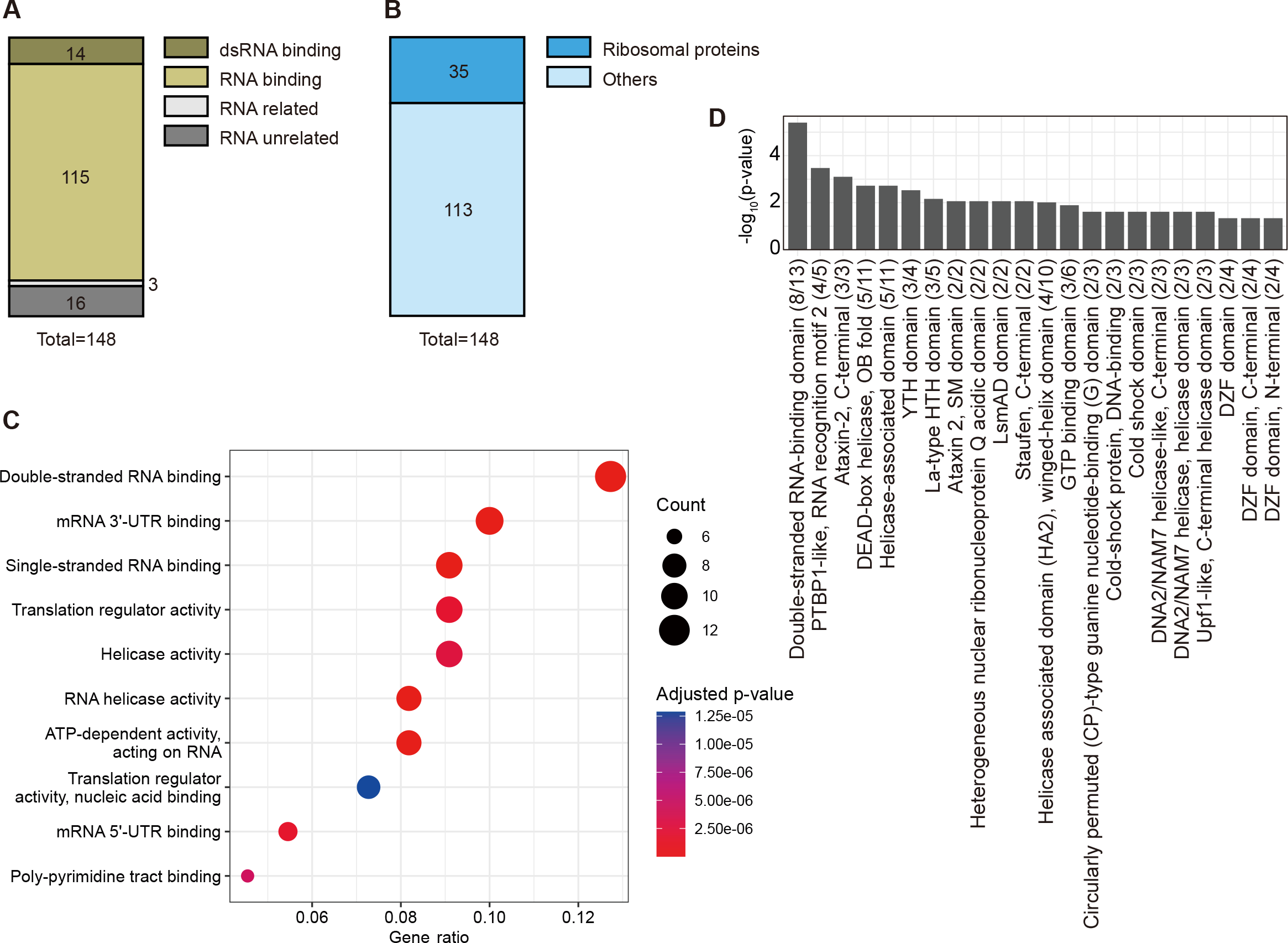
Characterization of K1 captured proteins. (A) Categorization of identified candidate dsRBPs based on GO terms in molecular function. (B) Number of ribosomal subunit proteins in the identified list. (C) A dot plot illustrating the enriched GO terms in molecular function among 113 candidate dsRBPs denoted as K1 interactome, excluding ribosomal subunits, analyzed using the clusterProfiler R package. (D) Significantly enriched protein domains and their respective p-values are shown. Numbers in the parentheses correspond to the number of proteins with the domain in the K1 interactome compared to the whole proteome background.

Enrichment analysis with GO of the molecular function of the identified K1 interactome also revealed various RNA-related GO terms as top hits, including ‘mRNA 3’-UTR binding’ and ‘Single-stranded RNA binding’. Notably, the GO term ‘Double-stranded RNA binding’ was the most significantly enriched term (Figure 2C). Domain enrichment analysis based on InterPro domain annotations revealed significant enrichment of RNA binding domains and helicase domains (Figure 2D). In addition, proteins containing well-known RNA-interacting domains, such as ‘PTBP1-like, RNA recognition motif 2’, ‘YTH domain’, and ‘La-type HTH domain’, were detected as well. In particular, ‘Double-stranded RNA-binding domain’ was the most significant enriched term. These results collectively confirmed that our K1 interactome reflects the dsRNA interactome, including previously unappreciated proteins.

### Validation of dsRNA binding capability of K1 interactome

Many of the cellular dsRNAs are transcribed as a part of host RNAs. For example, most SINE RNAs captured by the dsRNA antibody are embedded in introns and UTRs of protein-coding genes and are transcribed as a part of the host mRNA.^44^ Consequently, our K1 interactome may contain RBPs that were bound in the single-stranded region in the vicinity of the dsRNA region that was recognized by the K1 antibody. To exclude such a possibility and improve the accuracy of our putative dsRNA interactome, we performed two additional LC-MS/MS analyses (Figure 3A).

**Figure 3.**
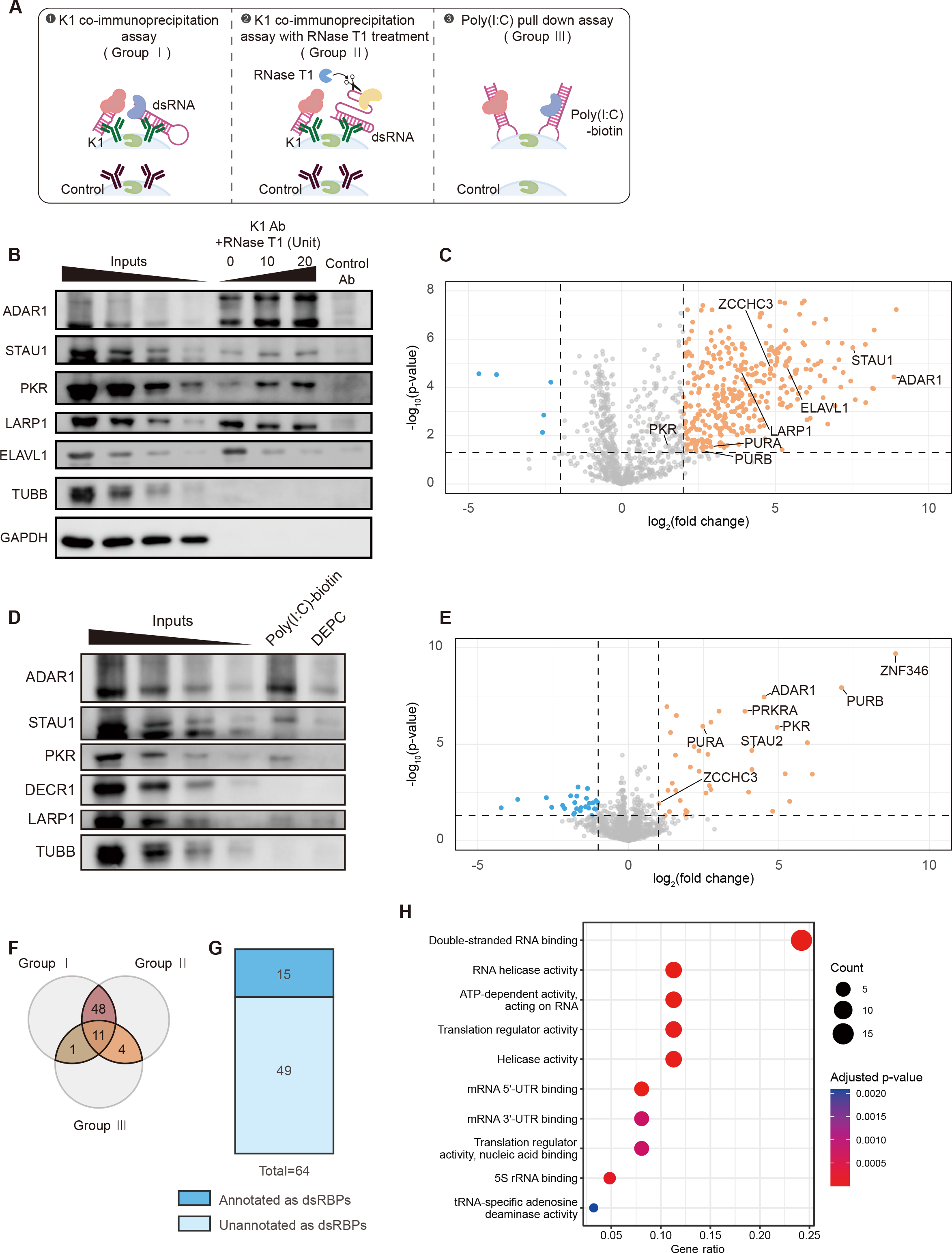
K1 capture with RNaseT1 treatment and poly(I:C) pull-down assay. (A) Schematics of two validation strategies using RNase T1 treatment in K1 IP and poly(I:C) pull-down followed by LC-MS/MS. (B) Western blot showing selected K1 captured proteins under RNase T1 treatment. (C) Volcano plot depicting the proteins captured by K1 with RNase T1 treatment, along with the mass spectrometry. Proteins that satisfy the filtering criteria with i) log_2_(fold change) ≥ 2 and ii) p-value < 0.05 are depicted in orange.Log_2_(fold change) was calculated as the difference between averaged log_2_(LFQ intensity) of 4 replicates in each experimental group. Statistical significances were calculated using two-sided Student’s t-tests. (D) Western blot of selected K1 captured proteins in poly(I:C) pull-down eluates. (E) Volcano plot showing poly(I:C) pull-down followed by LC-MS/MS results. Proteins that satisfy the filtering criteria with i) log_2_(fold change) ≥ 1 and ii) p-value < 0.05 are depicted in orange. Log_2_(fold change) was calculated as the difference between averaged log_2_(LFQ intensity) of 4 replicates in each experimental group. Statistical significances were calculated using two-sided Student’s t-tests. (F) Overlaps between the original K1 capture, RNase T1-treated K1 capture, and poly(I:C) pull-down proteins. (G) The number of previously annotated dsRBPs captured in our dsRNA interactome. (H) A dot plot illustrating the enriched GO terms in molecular function of the dsRNA interactome analyzed using the clusterProfiler R package.

First, we treated cell lysates with RNase T1, an endonuclease that cleaves after guanine residue of ssRNAs, during IP with the K1 antibody. The addition of the RNase T1 enzyme would ensure that RNA captured by the K1 antibody is fully double-stranded and remove contaminating ssRNA interactors. Feasibility test with K1 IP followed by western blotting showed that canonical dsRBPs, such as ADAR1, STAU1, and PKR, showed enhanced enrichment in RNase T1-treated samples (Figure 3B). La-related protein 1 (LARP1), one of the K1 interactome proteins, also retained moderate enrichment in RNase T1-treated samples. Although the degree of enrichment decreased, it is clearly higher than that of the animal-matched IgG control (Figure 3B). As a ssRNA-binding control, we used embryonic lethal, abnormal vision-like RNA binding protein 1 (ELAVL1) that recognizes AU-rich region in the 3’ UTR and controls RNA stability.^52^ Consistent with our expectation, RNase T1 treatment significantly reduced the enrichment of K1 co-IPed ELAVL1 (Figure 3B). As a negative control, we again used TUBB and GAPDH, which were undetected in any of the samples analyzed (Figure 3B).

We then prepared a large-scale RNase T1-treated K1 co-IPed sample for LC-MS/MS analysis. We performed LC-MS/MS analysis on four biological replicates, and statistical analysis showed good correlations between the replicates (Figure S2A). Of note, the correlation values between different experimental groups were less than those between K1 experimental and control groups (Figure S2A). This trend was consistent with the PCA plot showing clear clustering (Figure S2B). A total of 1,096 proteins were detected, and notably, RNase T1 treatment yielded a significantly higher proportion of proteins captured by K1 antibody compared to the animal-matched control (Figure 3C). This result reflects an improved degree of enrichment for dsRBPs, as shown by the western blotting results of ADAR1, STAU1, and PKR when RNase T1 was treated (Figure 3B). One possibility is that RNase T1 treatment reduced the ssRNA contaminants and short-structured RNAs that are recognized by the K1 antibody, hence resulting in improved signal-to-noise ratio for dsRBPs that interact with RNAs with extended stretch of double-stranded regions. As the RNase T1-treated K1 interactome contained too many hits, we used more stringent criteria of p-value less than 0.05 and a fold change of greater than four, which yielded 348 proteins. After excluding 76 ribosomal proteins, the GO analysis of the remaining 272 proteins annotated as RNase T1-treated K1 interactome revealed a significant enrichment of helicase-related GO terms, with the dsRNA binding GO term ranking fourth (Figures S2C, S2D, and Table S2).

Second, we utilized synthetic dsRNAs, poly(I:C), to compare the poly(I:C)-interacting proteins with the K1 interactome (Figure 3A). Prior to LC-MS/MS analysis, we performed IP with streptavidin-coated magnetic beads after incubating cell lysates with biotinylated poly(I:C) followed by western blotting. Well-known dsRNA-binding proteins (ADAR1, STAU1, and PKR) again showed good enrichment compared to control without poly(I:C) while TUBB did not show any enrichment (Figure 3D). Interestingly, K1 interactome proteins exhibited quite variable results. DECR1 was no longer detected in poly(I:C) pulled-down samples while LARP1 showed a moderate affinity for poly(I:C) (Figure 3D).

Mass spectrometry analysis of the poly(I:C) bound proteome revealed high enrichment of several dsRBPs, such as ADAR1, PKR, PRKRA, and STAU2, and showed a good correlation between biological replicates and distinguishable clusters in PCA analysis (Figures 3E, S3A, and S3B). In addition, proteins such as purine-rich element binding protein A (PURA), purine-rich element binding protein B (PURB), and zinc finger CCHC-type containing 3 (ZCCHC3) showed strong enrichment in all three types of LC-MS/MS analyses (Figures 1D, 3C, 3E, and S4). To delineate the poly(I:C) interactome, we identified 38 proteins meeting the criteria of p-value less than 0.05 and a fold change greater than two. Excluding one ribosomal protein, 37 proteins were designated as constituents of the poly(I:C) interactome, of which 12 were well-documented dsRBPs (Figure S3C and Table S3). The GO enrichment analysis highlighted dsRNA binding as the most significantly enriched term within this poly(I:C) interactome (Figure S3D).

The interactomes obtained through the aforementioned three methods represent proteins capable of binding to different types of dsRNA ligands, albeit sharing a common potential for dsRNA binding. As a result, in the process of discovering potential dsRNA interactomes without restricting the dsRNA ligand to poly(I:C), we selected proteins present in the interactomes in at least two of the three methods and denoted them as “dsRNA interactome” (Figures 3F and S4). A total of 64 proteins, including 15 previously annotated dsRBPs, were obtained (Figure 3G), and these proteins were further analyzed for their potential functional role, as shown below. Indeed, the GO analysis of these 64 proteins showed typical features of dsRBPs, such as ‘Double-stranded RNA binding’ (Figure 3H).

### Putative dsRBPs regulate IFN response to poly(I:C) and replication of the HCoV-OC43 virus

Currently, the most well-characterized function of dsRNA interacting proteins is the regulation of innate immune response, such as type I IFN response, during exogenous RNA signaling. For instance, RIG-I and MDA5 serve as cytosolic dsRNA sensors that trigger RLR-mediated IFN-β response,^53^ while ADAR1 disrupts dsRNAs with A-to-I editing to suppress autoimmune response to self-RNAs.^28^ Thus, we envisioned that many of the newly identified dsRBP candidates might also regulate the type I IFN response when exogenous RNA was introduced. To test, we carried out a CRISPR-Cas9-based loss-of-function screening in response to poly(I:C) transfection using secreted IFN-β level and cell viability as readouts (Figure 4A). Of note, we switched our experimental system to A549 lung adenocarcinoma cells because HEK293T cells showed too weak response to poly(I:C) transfection (Figures S5A and S5B).

**Figure 4.**
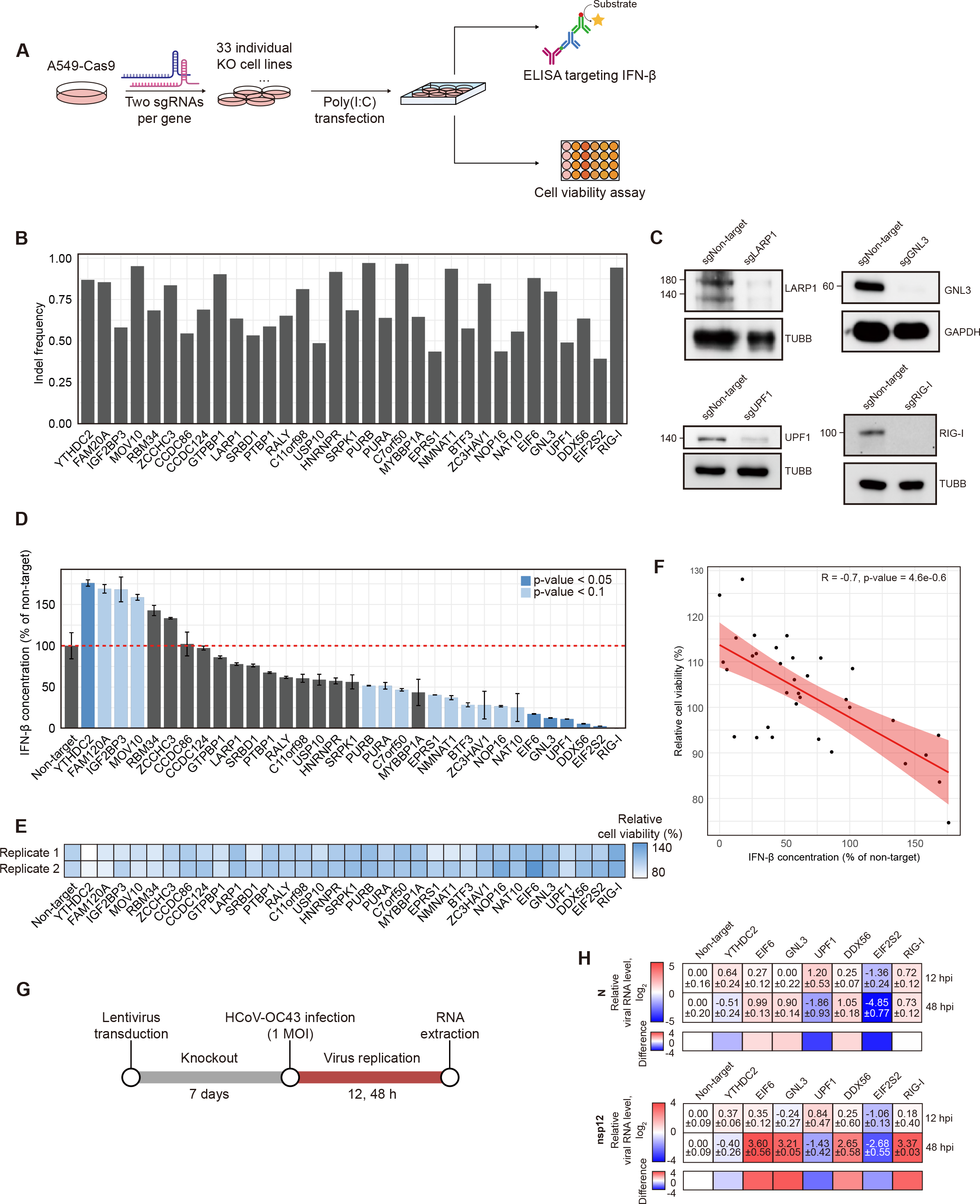
The immunoregulatory potential of dsRNA interactome. (A) Schematics of the CRISPR screening procedures. Each gene in the dsRNA interactome was targeted with a mixture of two sgRNAs. The KO cells were treated with poly(I:C), and secreted IFN-β level and cell viability were analyzed as readouts. (B and C) Validation of the KO efficiency with high throughput target DNA sequencing (B) and western blotting of selected proteins (C). (D) Normalized secreted IFN-β levels in the dsRNA interactome KO cells compared to NT control. Genes whose KO resulted in a significant change in IFN-β expression are shown as dark blue. An average of two biological replicates are shown with error bars denoting s.e.m. Statistical significance was calculated using a two-sided Student’s t-test, and p-values are indicated by the color of the bar. (E) The effect of poly(I:C) transfection in the indicated KO cells on cell viability, normalized against that of the NT cells. (F) Pearson correlation between IFN-β level and cell viability. (G) Schematic HCoV-OC43 infection followed by CRISPR KO in A549-Cas9 cells. (H) Change in N sgRNA and nsp12 gRNA levels in indicated KO cells 12 and 48 hpi. An average of three biological replicates are shown with s.e.m indicated as error bars. Below is the log_2_ difference in viral gene expression between 12 and 48 hpi.

Two sgRNAs were designed and prepared for the KO of each of the 32 genes curated from the dsRNA interactome. Of note, we selected 32 proteins out of 64 based on their reported molecular function. We first validated the experimental robustness by employing the well-known cytosolic dsRNA sensor, RIG-I, as a positive control. Three non-target (NT) sgRNAs that have no matching sequence in the entire human genome were designed to serve as a negative control. The high indel frequency at the genome level and the reduction in protein expression at the protein level in each generated KO cell confirmed that employing two sgRNAs per gene is sufficient to draw near complete removal of the targets (Figures 4B and 4C). Of note, we checked the protein expression of UPF1 as UPF1 showed one of the lowest indel frequencies of 49%, and our western results showed a near-complete reduction in the protein expression (Figure 4C).

We then transfected 1 ng/ml of poly(I:C) and measured secreted IFN-β levels from individual KO cells via ELISA assay. Tolerance towards the cytotoxic effect of poly(I:C) was also assessed by measuring the remaining cell population with CCK8 assay (Figures 4D and 4E). We found that KO of most genes resulted in significantly reduced levels of IFN-β compared to NT control (Figure 4D). One notable gene was zinc finger CCCH-type antiviral 1 protein (ZC3HAV1), which has been shown to suppress translation and facilitate the degradation of viral mRNA.^54–57^ Inhibition of ZC3HAV1 expression following influenza virus infection resulted in the suppression of IFN-β production, thereby promoting viral replication.^58^ Similarly, our screening result also showed decreased IFN-β production during antiviral response in ZC3HAV1 KO cells. On the contrary, in cells where YTH N^6^-methyladenosine RNA binding protein C2 (YTHDC2), family with sequence similarity 120 member A (FAM120A), insulin-like growth factor 2 mRNA binding protein 3 (IGF2BP3), and Mov10 RNA helicase (MOV10) KO, the release of IFN-β in response to poly(I:C) was enhanced. In particular, YTHDC2 and IGF2BP3 are reader proteins that recognize N^6^-methyladenosine (m^6^A)-modified RNAs, influencing the fate of mRNA and RNA metabolism. Previous studies have indicated that disturbances in the m^6^A machinery result in changes in the spread of various viruses due to irregular methylation of viral RNA,^59–61^ and it is speculated that the m^6^A readers, YTHDC2 and IGF2BP3, may be involved in this response. Furthermore, we observed that cells exhibiting hyper-activated IFN-β response were more susceptible to poly(I:C)-induced cell death, whereas cells with suppressed IFN-β response showed relatively less occurrence of cell death (Figure 4E). In other words, the IFN-β concentration in response to poly(I:C) and the cell survival rate induced by poly(I:C) showed a negative correlation (Figure 4F).

We further investigated the potential anti-or pro-viral roles of the putative dsRBPs. We focused on the genes that showed significant changes in secreted IFN-β levels with p-value < 0.05 and examined the effect of knocking out these genes in the replication of HCoV-OC43 (Figure 4G). HCoV-OC43 is a positive-sense, single-stranded RNA virus that generates dsRNA intermediates during its replication and transcription processes.^62,63^ As a positive control, we used RIG-I, whose KO resulted in enhanced viral replication. We found that eukaryotic translation initiation factor 6 (EIF6), G protein nucleolar 3 (GNL3), and DEAD-box helicase 56 (DDX56) KO cells exhibited an increase in viral replication over time, which is consistent with their effect of decreased IFN-β levels (Figure 4H). On the other hand, KO of YTHDC2, which exhibited an activated IFN-β response to poly(I:C), decreased the viral replication (Figure 4H). Of note, in YTHDC2 KO cells, the initial viral replication was facilitated when we analyzed 12 hours post-infection (hpi), but the replication was clearly reduced 48 hpi (Figures 4H, S5C, and S5D).

## DISCUSSION

In this study, we report a systematic and comprehensive effort to elucidate a repertoire of proteins that interact with cellular dsRNAs by directly capturing proteins bound to dsRNAs using K1 antibodies. Such method was employed to overcome the limitations of low UV crosslinking efficiency between dsRNA and dsRBP and to characterize dsRBPs that can interact with diverse classes of cellular dsRNAs rather than a single specific RNA bait. We further improved the reliability of our K1 interactome by utilizing RNase T1 to remove ssRNA-bound proteins and by employing poly(I:C) as a canonical dsRNA to capture interacting proteins. In the end, we discovered 64 putative dsRBPs, denoted as dsRNA interactome. We further investigated the functional role of selected dsRBP candidates via CRISPR-Cas9 KO screening with IFN-β and cell viability as readouts. Interestingly, we found that the KO of most dsRBPs resulted in decreased sensitivity to the dsRNA stressor poly(I:C), indicating that these proteins might function as mediators of innate immunity during exogenous RNA sensing. Indeed, we showed that cells deficient in these proteins are more vulnerable to HCoV-OC43 virus infection, underscoring the significance of dsRBP-mediated immune regulation.

Our study provides valuable resources and insights into the cellular dsRNA interactome. For instance, two of the proteins whose KO enhanced IFN-β levels to poly(I:C) stimulation were m^6^A readers, YTHDC2 and IGF2BP3. Previously, studies showed the potential role of m^6^A modification on innate immunity where m^6^A writer methyltransferase 3 (METTL3) induces RNA methylation to prevent dsRNA formation while m^6^A reader YTH N^6^-methyladenosine RNA binding protein F2 (YTHDF2) recognizes m^6^A modified dsRNAs to facilitate their degradation.^64–67^ Our data support that m^6^A modification may occur on dsRNAs as m^6^A readers are captured by the K1 antibody. In addition, the increased IFN-β levels to poly(I:C) stress in cells deficient in YTHDC2 and IGF2BP3 further suggest that the removal of m^6^A-modified RNAs by these reader proteins can affect innate immunity during exogenous RNA sensing. Of note, since poly(I:C) cannot be methylated, our results indicate that basal dsRNA expression may affect the sensitivity of exogenous RNA sensing and that m^6^A modification may regulate the basal dsRNA level. In addition, considering that METTL3 can directly modify viral RNAs, the removal of m^6^A-modified dsRNAs by the readers might be an evolved viral defense mechanism.^68^ Interestingly, the KO of YTHDC2 resulted in the initial enhancement of HCoV-OC43 viral replication that is eventually suppressed over time. Further investigation is required to delineate the potential selectivity of m^6^A modification on dsRNAs and its role in innate immunity to cellular dsRNAs.

Our dsRNA interactome contains many proteins that bind to 3′ UTRs. A recent report showed that UPF1 interacts with Ras-GTPase-activating protein binding protein 1 (G3BP1) in highly structured 3′ UTRs, leading to structure-mediated RNA decay,^69^ which aligns with our findings. It is noteworthy that the depletion of nonsense-mediated decay components, including UPF1, reduced IFN-β release upon poly(I:C) stimulation, concomitant with an escalation in viral replication at 12 hpi. This result aligns with the tendency of UPF1 to recognize and degrade incoming viral RNA genomes shortly after the onset of infection.^70,71^ In addition to UPF1, our list contains G3BP2 and LSM12, two components of stress granules.^72,73^ Notably, our pull-down experiments were conducted in HEK293T cells without any external stress, under which no stress granules were formed. Yet, our data clearly suggest that stress granule components are bound to cellular dsRNAs. This result is consistent with a recent study that showed aged mRNAs are bound by stress granule proteins even without granule formation.^74^ In the light of our study, cellular dsRNAs may work as seed molecules to facilitate liquid-liquid phase transition during stress conditions. Such possibilities should be investigated in detail.

Lastly, we found many proteins that were previously shown to directly interact with viral RNAs and regulate their replication. A previous study showed that ZCCHC3 binds to viral RNAs and facilitates the recognition of these RNAs by RIG-I-like receptors.^75^ Our finding suggests that ZCCHC3 may also bind to cellular dsRNAs and regulate their recognition by dsRNA sensors to regulate the downstream IFN response. In addition, we were surprised to see LARP1 on our list. Recent studies showed that LARP1 directly binds to SARS-CoV-2 RNA and regulates the downstream immune response to these viral RNAs.^38,39^ Based on our results, it is possible that LARP1 recognizes dsRNAs or dsRNA-like structured RNAs and promotes their recognition by dsRNA sensors. Therefore, our study provides valuable insights into the potential mode of interaction between RBPs that can recognize viral RNAs to mediate cellular defense against infection.

Overall, our work highlights the existence of several previously unappreciated potential dsRBPs that play central roles in modulating the antiviral response during exogenous RNA sensing as well as during defense against viral infection. We provide valuable resources of putative dsRBPs for further investigation, and our approach can be further extended to various physiological and pathological contexts to broaden our understanding of dsRNA biology.

### Limitations of the study

While this extensive dataset provides interesting insights into the dsRNA interactome, the study is subject to several limitations that necessitate further investigation. Our study was mainly based on dsRNA capture using the K1 antibody and could not provide specific interacting dsRNA species for individual dsRBP candidates. Consequently, our results might contain pseudo-dsRBPs that were inadvertently co-eluted with protein complexes formed via protein-protein interactions. For example, our list contains many proteins involved in stress granule formation. Although many of these proteins are shown to have RNA-binding properties, their direct interaction with dsRNAs must be examined individually before concluding that these proteins are indeed dsRBPs. In addition, although the majority of cellular dsRNAs and the RNAs recognized by the dsRNA antibody are derived from SINEs,^44^ the class of cellular dsRNAs that may interact with our dsRNA interactome is unclear. Considering the biophysical nature of dsRNA-RBP interaction that RBPs recognize the structural features of the dsRNA due to the narrow major groove of the A-form helix of the dsRNA,^5,48^ it is likely that our putative dsRBPs will bind to SINE dsRNAs. However, due to subcellular localization or liquid-liquid phase transition, certain dsRBPs will preferentially bind to a selective class of dsRNAs. The dsRNA substrates of the identified putative dsRBPs need to be discovered through methods such as formaldehyde crosslinking immunoprecipitation and sequencing. Lastly, the K1 antibody can detect proteins binding to dsRNA molecules with helices longer than 40 bp, making K1 insufficient to capture proteins binding to short dsRNAs.^43,76^ Particularly, these short dsRNAs can serve as agonists for RIG-I.^77,78^ Moreover, for dsRNAs to be recognized by K1, proteins must not cover the entire dsRNA and exclude regions to allow the K1 antibody to bind on the same RNA molecule. Perhaps, these are the reasons why our list lacks RIG-I and MDA5, as the latter oligomerizes along the dsRNA and may occupy the entire dsRNA.^79,80^ Despite these limitations, our study successfully identified cellular dsRNA interactome and explored their roles in innate immunity during exogenous RNA sensing and viral infection.

## Supporting information

Key resources table

## Acknowledgments

We thank all members of the Jong-Seo Kim, Sangsu Bae, and Yoosik Kim laboratory for helpful discussion and comments on the paper. We acknowledge the KAIST Analysis Center for Research Advancement for experimental equipment. This study was supported by the Basic Research Laboratory Program through the National Research Foundation of Korea (NRF-2021R1A4A3032789). J.-S.K. was supported by the Institute for Basic Science of the Ministry of Science and ICT of Korea (IBS-R008-D1).

## Author contributions

J.L., N.L., S.J., and Y.K. designed the study and analysis. J.L., N.L., S.J., J.K., S.M., M.J., and S.-H.L. performed experiments. Y.-k.L. set up the virus experiment while S.K. assisted in establishing the CRISPR screening. J.K. assisted in establishing the K1 co-IP experiment. J.-S.K. and Y.K. supervised the study. J.L., N.L., and Y.K. wrote the manuscript with contributions from S.J. and S.-H.L. All of the authors subsequently reviewed and edited the manuscript.

## STAR★METHODS

### RESOURCE AVAILABILITY

#### Lead contact

Further information and requests for resources and reagents should be directed to and will be fulfilled by the lead contact, Yoosik Kim.

#### Materials availability

All reagents and resources used for this study are available upon request to the lead contact.

#### Data and code availability

- Original western blot images will be deposited at Mendeley. The DOI will be listed in the key resources table upon acceptance of this manuscript. All LC-MS/MS data utilized in this study can be accessed from the PRIDE database under accession number: PXD053100.
- This paper does not report the original code.
- Any additional information required to reanalyze the data reported in this paper is available from the lead contact upon request.

### EXPERIMENTAL MODEL AND SUBJECT DETAILS

#### Cell lines

All 5 cell lines used in this study (HEK293T, A549, A549-Cas9, RD, and HCT-8) were maintained in culture media supplemented with 10% FBS and routinely cultured at 37°C with 5% CO_2_. FBS were thawed in water at room temperature (RT) and were not heat-inactivated. RPMI 1640 (Welgene, LM011-01) were used when culturing HCT-8 while DMEM with high glucose (Welgene, LM001-05) were used for HEK293T, A549, A549-Cas9, and RD cells. Cells were frequently tested negative for mycoplasma contamination.

### METHODS DETAILS

#### K1 capture

HEK293T cells were cultured and harvested with a scraper. The collected cell pellet was washed twice with cold DPBS. The cell pellet was then lysed through a syringe with lysis buffer (TD buffer supplemented with 1% NP40) supplemented with protease inhibitor cocktail and recombinant RNase inhibitor to prevent any denaturation of proteins and RNAs during K1 co-IP. Before incubating the magnetic protein A beads with lysate, 50 μl of the beads were washed once with 300 μl of lysis buffer and then incubated with 10 μg of the K1 antibody (Scicons) or an animal-matched control antibody for 3 h at 4°C with rotation. 50 μl of antibody-coated magnetic protein A beads were washed twice with lysis buffer and incubated with 300 μl of the lysate for 3 h at 4°C with rotation. The beads were washed three times with lysis buffer and then centrifuged. Then, 5X SDS-PAGE loading buffer was added to the appropriate volume, and the sample was denatured at 95°C for 10 min. The samples were then analyzed with western blots.

After the final wash step of the protein A beads using a wash buffer lacking NP40, 8 M urea in 50 mM ABC (ammonium bicarbonate) buffer was added to the centrifuged beads for LC-MS/MS analysis. The sample was then incubated at RT overnight with shaking in a thermomixer at 600 rpm (Eppendorf). The sample was centrifuged, and the supernatant was analyzed for LC-MS/MS.

#### Protein digestion for LC-MS/MS analysis

The eluted proteins in 8 M urea in 50 mM ABC buffer were denatured by incubating for 1 h at 37°C in a thermomixer (Eppendorf). The samples were treated with dithiothreitol at a concentration of 10 mM and followed by 40 mM iodoacetamide in the dark, each incubated for 1 h at 37°C. To lower the urea concentration below 1 M, the samples were further diluted with 50 mM ABC buffer. Subsequently, CaCl_2_ was added to make the final concentration of 1 mM. Sample digestion was carried out with MS grade trypsin (Thermo Fisher Scientific) at a weight ratio of 1:50 (trypsin to protein), with an overnight incubation at 37°C. The digestion process was followed by SPE clean-up using a C18 cartridge (Supelco). The digested peptides were dried using a SpeedVac (Thermo Fisher Scientific) and then reconstituted in 50 mM ABC for LC-MS/MS analysis.

#### LC-MS/MS analysis

In-house packing of both analytical capillary columns (100 cm×75 μm i.d.) and trap columns (2 cm×150 μm i.d.) utilized 3 μm Jupiter C18 particles (Phenomenex). A consistent column temperature of 45°C for the long analytical column was achieved through the use of a dedicated column heater (Analytical Sales and Services). Chromatographic elution was performed using a NanoAcquity UPLC system (Waters) at a flow rate of 300 nl/min over a total run time of 150 min, which included a linear gradient over 100 min, transitioning from 95% solvent A (0.1% formic acid in water) to 40% solvent B (0.1% formic acid in acetonitrile). Two mass spectrometers were employed for LC-MS/MS analysis, both equipped with a specialized nanoelectrospray ion source designed in-house: the Orbitrap Eclipse and the Orbitrap Fusion Lumos (Thermo Fisher Scientific). The setup for the Orbitrap Eclipse involved acquiring precursor ions within an m/z range of 300–1500 at a resolution of 120K. Isolation of the precursor for MS/MS analysis was performed using a 1.4 Th. High-energy collisional dissociation (HCD) for peptide sequencing was applied at an energy setting of 30%. MS2 scans were performed at a resolution of 30K, targeting an AGC of 1 × 10⁵ with a maximum ion injection time (ITmax) of 54 ms. For the Orbitrap Fusion Lumos, the settings were an m/z range of 300–1800 at a resolution of 60K, MS2 scans at a resolution of 7.5K, an AGC target of 5 × 10⁴, and an ITmax of 22 ms. The mass spectrometry proteomics data have been deposited to the ProteomeXchange Consortium via the PRIDE^83^ partner repository with the dataset identifier PXD053100 and 10.6019/PXD053100.

#### RNase T1-treated K1 co-IP and poly(I:C)-biotin pull-down

For the RNase T1-treated K1 co-IP experiment, HEK293T cells were harvested with ice-cold lysis buffer with 1% NP40. Lysate was prepared with syringe lysis at 4°C with a minimum 100 passing through the syringe. A protease inhibitor cocktail was added to prevent any denaturation of protein. Magnetic protein A beads (Bioneer) were prepared by washing once with the lysis buffer. Beads were then incubated with a K1 antibody (Scicons) or an animal-matched control antibody for 3 h at 4°C. After incubation, the beads were washed three times with lysis buffer. The prepared lysate was added to the beads together with an 20 unit of RNase T1 to remove ssRNAs. Samples containing RNase T1 were incubated at 4°C for 3 h with rotation for binding and washed three times with lysis buffer. Additional wash steps using wash buffer without NP40 was carried out twice to remove any residual detergents. Samples were eluted with the elution buffer (8M Urea in 50 mM ABC buffer) with rotation at 37°C.

For poly(I:C)-biotin pull-down, HEK293T lysate collected using ice-cold lysis buffer containing 1% NP40 supplemented with protease inhibitor cocktail was prepared by sonication followed by centrifugation for 10 min at 4°C. Streptavidin beads (Bioneer) were prepared by washing once with the lysis buffer. Beads were then incubated with biotin-poly(I:C) conjugates (Invitrogen) for 3 h at 4°C. After incubation, beads were washed three times with fresh lysis buffer. Prepared lysate was added to the beads, and samples were incubated at 4°C for 3 h for binding. After incubation, the beads were washed three times with lysis buffer, followed by two additional washes using wash buffer without NP40. Samples were eluted with elution buffer (8M Urea in 50mM ABC buffer) with rotation at 37°C.

Extracted samples from RNase T1-treated K1 co-IP and poly(I:C) pull-down methods were immediately frozen at −80°C until further analysis with LC-MS/MS. For the western blot assay, 5X SDS-PAGE loading buffer was added instead of the elution buffer, and the mixture was boiled at 95°C for 10 min.

#### Western blotting

For K1 antibody co-IPed samples, cell pellets were lysed with TD buffer supplemented with 1% NP40. The buffer was additionally supplemented with a protease inhibitor cocktail and RNase inhibitor. To validate the target protein expression in KO cells, lysates in target KO cells were prepared with the sonication using RIPA lysis buffer and then run through 10% SDS PAGE gel. The proteins on the gel were transferred to a PVDF membrane using the Amersham semidry transfer system. The membrane was blocked in 5% skim milk for 1 h at RT and incubated with primary antibody overnight at 4°C. The following primary antibodies were used: ADAR1(14175), PKR (12297), TUBB (2128), GAPDH (5174), ELAVL1 (12582), TUBB (86298), and RIG-I (3743) were purchased from Cell Signaling Technology; GAPDH (sc-32233) was purchased from Santa Cruz Biotechnology; LARP1 (A302-087A) and LARP4 (A303-900A) were purchased from Bethyl Laboratories; STAU1 (ab73478) and GNL3 (ab70346) were purchased from Abcam; DECR1 (PA5-80549) was purchased from Invitrogen; UPF1 antibody was provided by Yoon Ki Kim’s laboratory at KAIST. All antibodies were used at 1:1000 dilution.

#### Generation of dsRNA interactome KO cell lines

Cas9-BLAST and lentiGuide-Puro inserted sgRNA plasmids were designed and packaged into lentiviral vector using psPAX2 and pMD2.G, according to the protocol described in O.Shalem et al.^84^ To delete the target gene expression, two sgRNAs were designed and a mixture of two kinds of lentiviruses containing each sgRNAs was transduced into Cas9-expressing A549 cells, supplemented with 5 μg/ml polybrene to enhance transduction. The sequences of sgRNAs are listed in Table S4. 24 h post-transduction, cells were selected with puromycin at 5 μg/ml concentration, and the media was refreshed every other day. 7 days post-transduction, the KO efficiency was confirmed via high-throughput DNA sequencing and western blotting.

#### Poly(I:C) transfection

For poly(I:C) transfection, cells were seeded at a density of 4×10^5^ cells per well in a 6-well plate and allowed to stabilize in media without puromycin for one day. Poly(I:C) was transfected at 1 ng/ml using Lipofectamine 3000, p3000, and Opti-MEM media according to the manufacturer’s protocol. 16 h post-transfection, media was collected to detect the concentration of IFN-β via ELISA. Cell viability was also determined 16 h after poly(I:C) transfection.

#### High-throughput DNA sequencing

The cell pellet was resuspended in 100 μl proteinase K extraction buffer (40 mM Tris-HCl (pH8.0), 1% Tween-20, 0.2 mM EDTA, 2 mg proteinase K, 0.2% NP-40 (VWR)). The resuspended pellet was incubated at 60°C for 15 min, followed by 98°C for 5 min. PCR amplification was then carried out with KOD-Multi & Epi-(TOYOBO) according to the manufacturer’s protocol using 2 μl of genomic DNA as a template. The 1 μl of PCR product was subjected to next PCR amplification using primers containing next-generation sequencing adaptors. The primer sequences used for PCR are listed in Table S4. The PCR products were purified using a PCR purification kit (GeneAll) and then analyzed using an Illumina MiniSeq instrument. The sequencing results were analyzed using Cas-Analyzer software (http://www.rgenome.net/cas-analyzer/).^85^

#### ELISA and cell viability assay

For the ELISA experiment, cell culture media was collected and centrifuged at 4°C at 15,000 rpm for 30 min. The supernatant of the centrifuged sample was collected and used for ELISA assay. IFN-β human ELISA Kit (Thermo Fisher Scientific; 414101) was used following the manufacturer’s instructions to quantify IFN-β. 50 μl of sample diluent and 50 μl of the supernatant were added to each well of a 96-well plate and incubated for 1 h. The plate was aspirated and washed 3 times with wash buffer. The plate was exposed to a 100 μl of diluted antibody solution for 1 h and washed 3 times with wash buffer. 100 μl of HRP solution was added and incubated for 1 h. The plate was washed 3 times with wash buffer, and 100 μl of warmed TMB substrate solution was added. The plate was kept at RT in the dark for 15 min. After the incubation, 100 μl of stop solution was added, and the absorbance was measured at 450 nm using a microplate reader. Cell viability was determined using the Cell Counting Kit-8 (CCK-8) assay according to the manufacturer’s instructions. Specifically, 25 μl of CCK-8 solution was added to 250 μl media in each well of a 48-well plate, followed by incubation for 3 h at 37°C. The absorbance at 450 nm was then measured using a microplate reader.

#### Virus propagation and infection

For HCoV-OC43 propagation, 5 x 10^6^ HCT-8 cells were seeded into T-75 flasks and stabilized for 24 h. Cells were rinsed twice with warm DPBS and then exposed to 6 ml of virus-diluted media without FBS at a concentration of 0.5 MOI, as determined by plaque assay, for 1 h at 35°C. Following infection, the media containing the virus was substituted with reduced-serum media with 5% FBS, and cells were incubated at 35°C until harvesting. The titer of propagated HCoV-OC43 was measured using a 12-well plate with 5×10^5^ RD cells seeded. The virus solution, diluted by a factor of 10, was used to infect the cells for 1 h without FBS, followed by overlaying with 0.75% agarose containing reduced-serum media with 5% FBS. After 5 days, the number of plaques formed was determined by staining with crystal violet, counted, and the titer was calculated.

For HCoV-OC43 infection, 4×10^5^ cells were seeded in a 6-well plate and incubated with 5% CO_2_ at 37°C for one day. Cells were washed with warm DPBS twice and replaced with virus-diluted media without FBS as 1 MOI and incubated at 35°C for 1 h. Infected cells were washed one more time and replaced with reduced-serum media containing 5% FBS and incubated at 35°C for 48 h.

#### RNA purification and RT-qPCR

For total RNA extraction, 1 ml of TRIzol (Invitrogen) was added to the media-depleted cell monolayers. The TRIzol-treated sample was vortexed for 30 s after adding 200 μl of chloroform. The sample was centrifuged for 15 min at 4°C and 12,000 rpm, and 500 μl of supernatant was collected and added to a clean e-tube containing 30 μl NaOAc and 0.5 μl glycoblue. 500 μl of IPA was added, and the nucleic acids were precipitated overnight at −20°C. The sample was centrifuged at 15,000 rpm for at least 1 h at 4°C, and the precipitated nucleic acids were washed twice with 1 ml of 75% EtOH. To remove DNA, pellets were dissolved in 42 μl of TDW, and 2 μl DNase, 1 μl RNase inhibitor, and 5 μl DNase buffer were added. The sample was incubated at 37°C for 30 min. 150 μl of TDW and 200 μl of acid phenol were added, and the solution was vortexed for 30 s and centrifuged for 5 min at 12,000 rpm and RT. RNA was precipitated by adding 1 ml of 100% EtOH to the aqueous phase and incubating overnight at −80°C. After washing twice with 75% EtOH, the pellet was dried and dissolved with TDW. Following RNA isolation, 800 ng of RNA was reverse-transcribed using RevertAid transcriptase (Thermo Scientific) and random hexamers (Thermo Scientific). Subsequently, RT-qPCR was conducted using primer pairs specified in Table S4 and SYBR Green (Bioline), and the data were analyzed utilizing QuantStudio 1 (Thermo Scientific).

### QUANTIFICATION AND STATISTICAL ANALYSIS

#### Protein identification and statistical analysis of the dsRNA interactome

MS/MS data were analyzed using MaxQuant software (version 2.0.3.0) with the Andromeda search engine against the SwissProt Homo sapiens proteome database (version 2023.01, with 20,404 entries; Uniprot (http://www.uniprot.org/)). Mass tolerances of 10 ppm for precursor ions and 20 ppm for fragmentation ions were applied. The label-free quantitation (LFQ) and matching between runs were executed with specified search criteria: trypsin digestion, allowing up to 2 missed cleavages, fixed modification of carbamidomethylation at cysteine sites, and variable modifications of acetylation at protein N-termini, oxidation at methionine. The PERSEUS software was utilized as the statistical tool for identifying proteins within the dsRNA interactome and for quantitative assessment. LFQ intensities were log-transformed following the exclusion of reverse sequences and contamination. Protein groups reporting a minimum of two LFQ intensities under at least a single experimental group were included in subsequent analyses. Missing LFQ values were substituted using imputation from a normal distribution. A Student’s t-test, with false discovery rate (FDR) adjustments according to Benjamini-Hochberg at a 0.05 threshold, was employed for the statistical evaluation of protein groups. The same statistical methodology was used to investigate differentially captured protein groups within dsRNA interactomes.

#### Gene Ontology (GO) enrichment analysis

Based on the LC-MS/MS analysis results of K1 and animal-matched IgG co-IPed samples, proteins with a fold change over 2 and p-value below 0.05, excluding ribosomal subunits, were selected for GO analysis. For the proteomic results of RNaseT1-treated K1 capture, a more stringent criterion was applied, selecting proteins with a fold change of 4 or greater and a p-value of 0.05 or less, excluding ribosomal subunits, for GO analysis. To analyze proteomic results of poly(I:C) pulled-down samples for GO enrichment, proteins were selected with a fold change of 2 or greater and a p-value of 0.05 or less, excluding ribosomal subunits. Enrichment of GO analysis focusing on molecular function associated with enriched proteins in three kinds of dsRBP captures was conducted using a hypergeometric test in the clusterProfiler package (v4.4.4).^82^

#### Protein domain enrichment analysis

Protein domain annotations for all known human proteins were downloaded from InterPro.^86^ The protein domain profile of enriched proteins from LC-MS/MS was compared against that of background proteins, and enrichment and under-representation of each domain were calculated. The Fisher’s exact test was performed to calculate statistical significance, and p-values were adjusted for multiple testing with the Benjamini-Hochberg method.

### SUPPLEMENTAL INFORMATION

Document S1. Figures S1–S5.

Table S1. Excel file containing K1 interactome, related to Figures 2B, 2C, and 2D.

Table S2. Excel file containing RNase T1-treated K1 interactome, related to Figures S2C and S2D.

Table S3. Excel file containing poly(I:C) interactome, related to Figures S3C and S3D.

Table S4. Excel file containing oligonucleotide sequences, related to Figures 4B–4F, and 4H.

