## Supplementary material for "Cellular dsRNA interactome reveals the regulatory map of exogenous RNA sensing": Figures S1-S5

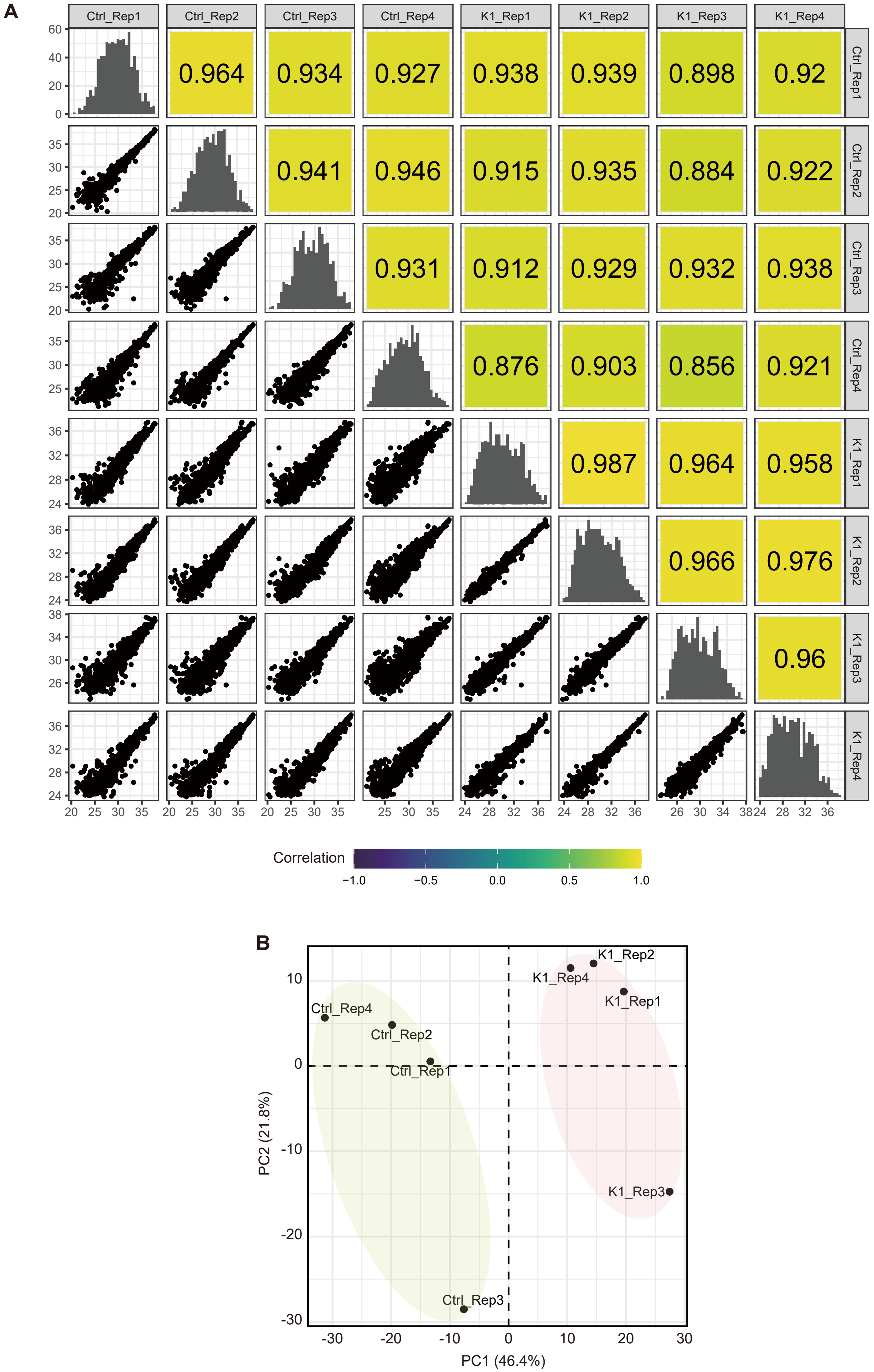
A
Ctrl_Rep1
Ctrl_Rep2
Ctrl_Rep3
Ctrl_Rep4
K1_Rep1
K1_Rep2
K1_Rep3
K1_Rep4
0.964
0.934
0.927
0.938
0.939
0.898
0.92
0.941
0.946
0.915
0.935
0.884
0.922
0.931
0.912
0.929
0.932
0.938
0.876
0.903
0.856
0.921
0.987
0.964
0.958
0.966
0.976
0.96
Correlation
−1.0
−0.5
0.0
0.5
1.0
B
K1_Rep2
K1_Rep4
K1_Rep1
K1_Rep3
Ctrl_Rep4
Ctrl_Rep2
Ctrl_Rep1
Ctrl_Rep3
PC2 (21.8%)
PC1 (46.4%)

**Figure S1. Mass spectrometry data analysis of K1 co-IPed samples, Related to Figure 1.** (A) Pairwise correlation matrix of proteins detected in each of four biological replicates of K1 co-IPed and control samples by LC-MS/MS. (B) PCA plot showing the distinct characteristics of each experimental group.

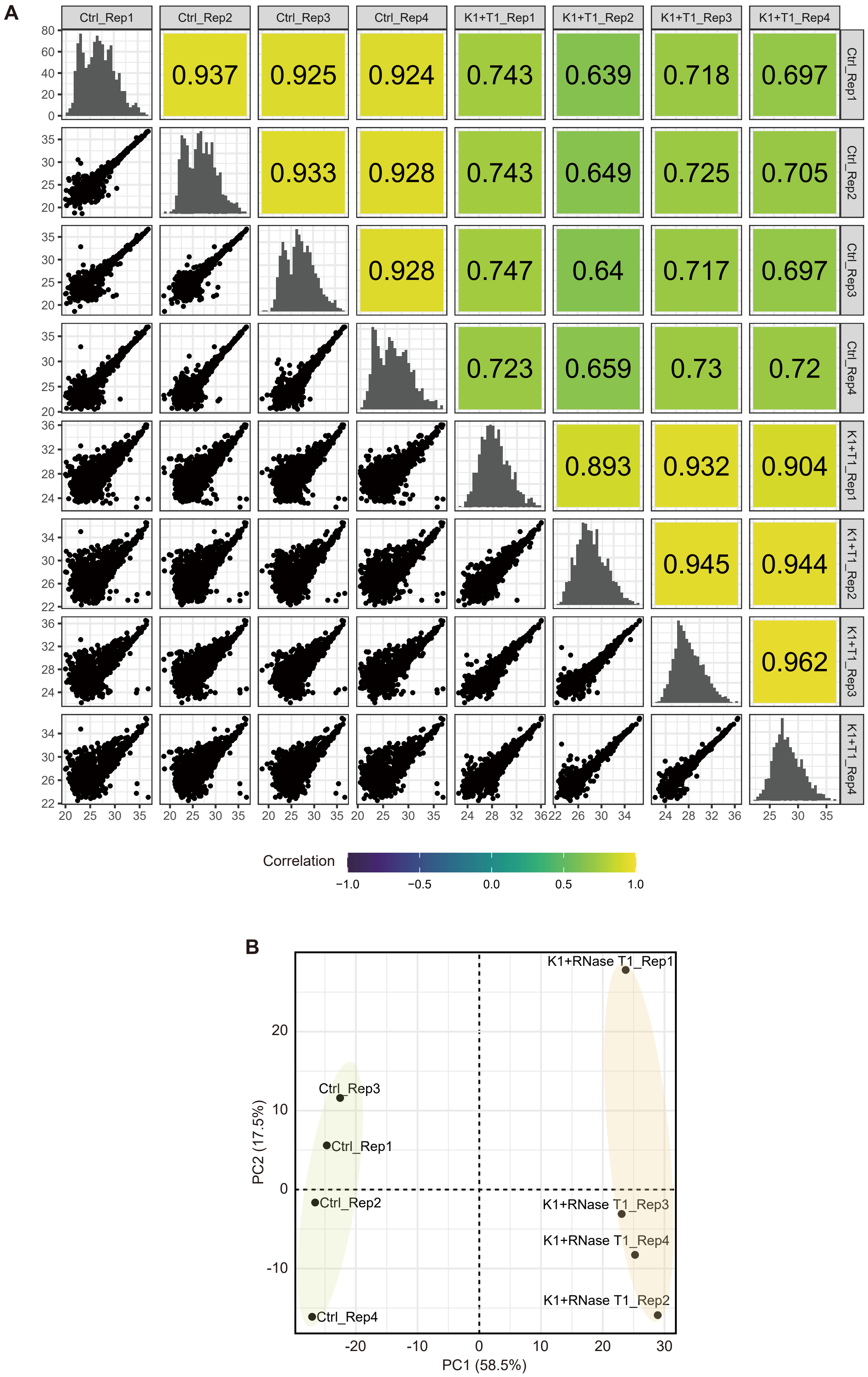
A
Ctrl_Rep1
Ctrl_Rep2
Ctrl_Rep3
Ctrl_Rep4
K1+T1_Rep1
K1+T1_Rep2
K1+T1_Rep3
K1+T1_Rep4
0.937
0.925
0.924
0.743
0.639
0.718
0.697
0.933
0.928
0.743
0.649
0.725
0.705
0.928
0.747
0.64
0.717
0.697
0.723
0.659
0.73
0.72
0.893
0.932
0.904
0.945
0.944
0.962
Correlation
−1.0
−0.5
0.0
0.5
1.0
B
K1+RNase T1_Rep1
Ctrl_Rep3
Ctrl_Rep1
Ctrl_Rep2
K1+RNase T1_Rep3
K1+RNase T1_Rep4
K1+RNase T1_Rep2
Ctrl_Rep4
PC2 (17.5%)
PC1 (58.5%)
20
10
0
-10
-20
-10
0
10
20
30

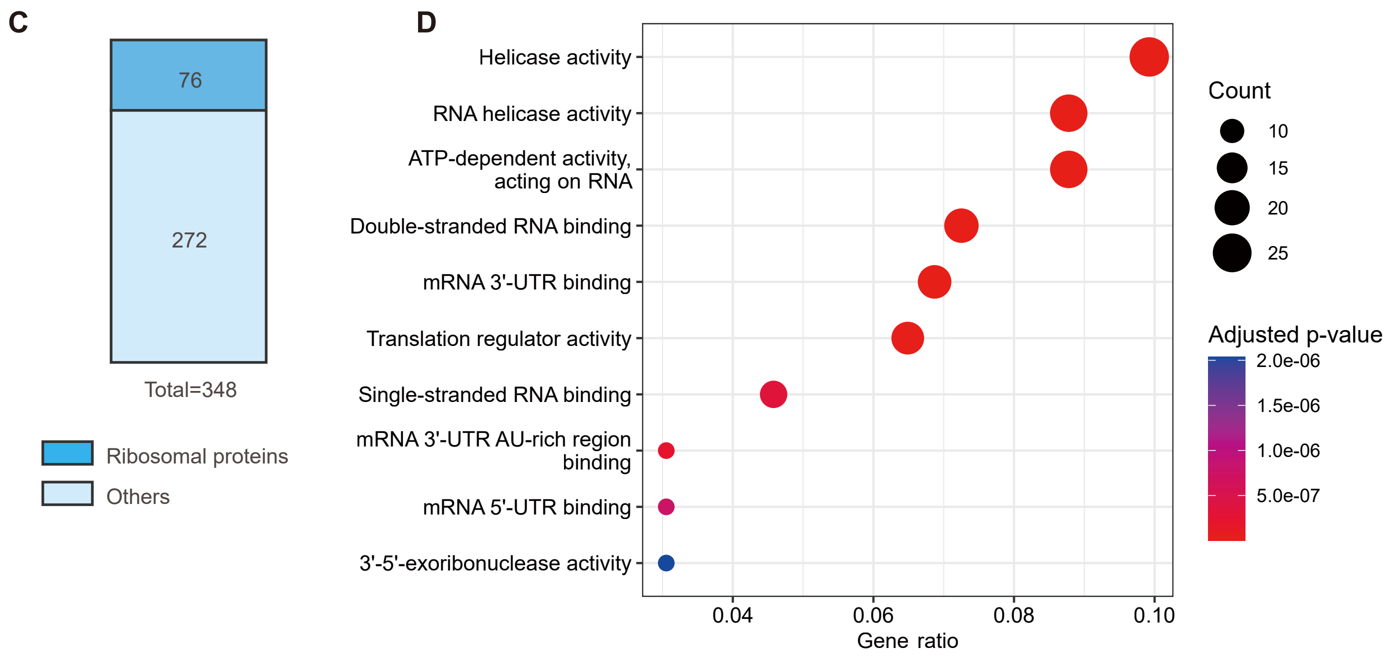


**Figure S2. Mass spectrometry data analysis of RNase T1-treated K1 co-IPed samples, Related to Figure 3.** (A) Pairwise correlation matrix of proteins detected in each of four biological replicates of RNase T1 treated K1 co-IPed and control samples by LC-MS/MS. (B) PCA plot showing the distinct characteristics of each experimental group. (C) Number of ribosomal and non-ribosomal proteins in the enriched proteins. (D) A dot plot illustrating the enriched GO terms in molecular function among RNaseT1-treated K1 interactome, analyzed using the clusterProfiler R package.

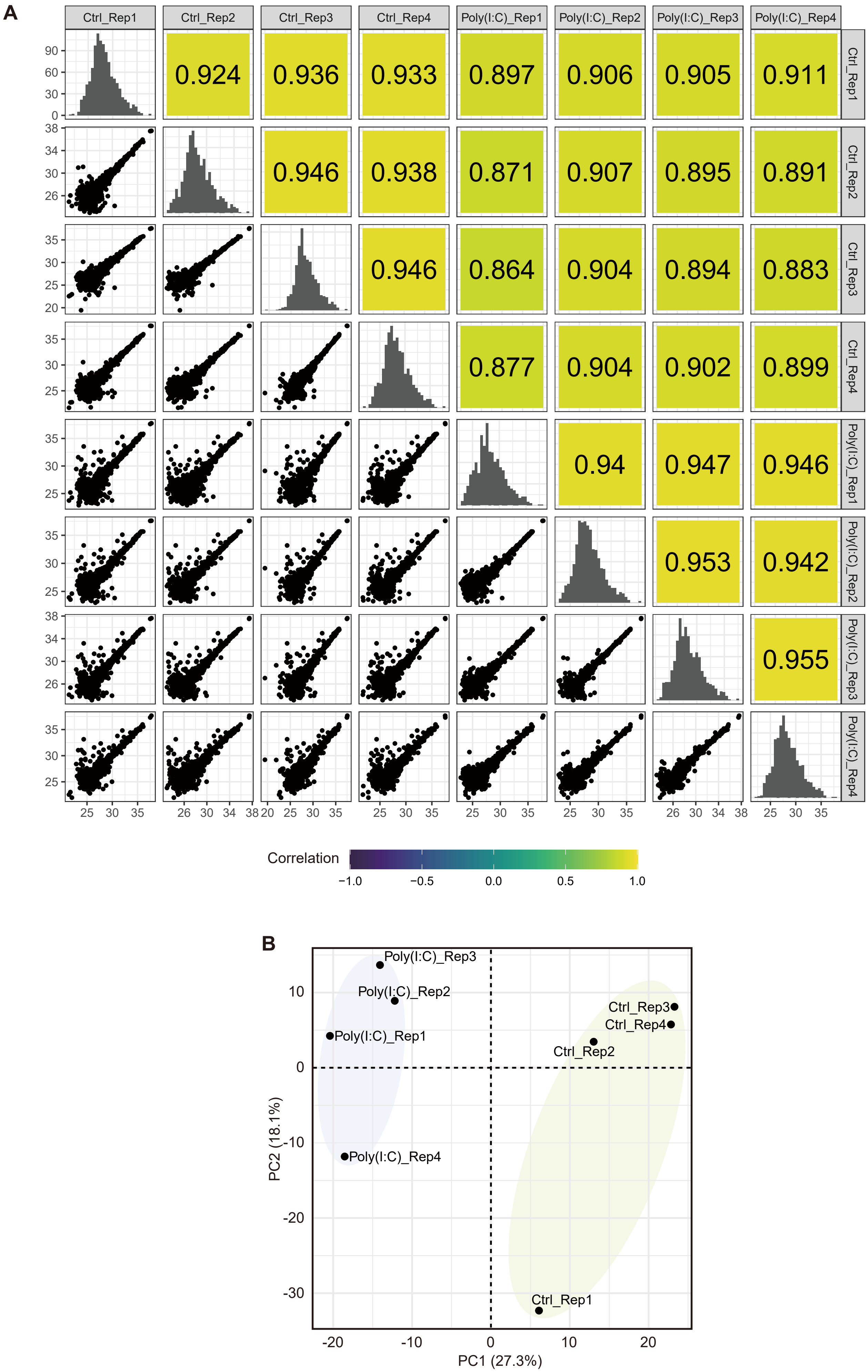
A
Ctrl_Rep1
Ctrl_Rep2
Ctrl_Rep3
Ctrl_Rep4
Poly(I:C)_Rep1
Poly(I:C)_Rep2
Poly(I:C)_Rep3
Poly(I:C)_Rep4
0.924
0.936
0.933
0.897
0.906
0.905
0.911
0.946
0.938
0.871
0.907
0.895
0.891
0.946
0.864
0.904
0.894
0.883
0.877
0.904
0.902
0.899
0.94
0.947
0.946
0.953
0.942
0.955
Correlation
−1.0
−0.5
0.0
0.5
1.0
B
Poly(I:C)_Rep3
Poly(I:C)_Rep2
Poly(I:C)_Rep1
Poly(I:C)_Rep4
Ctrl_Rep3
Ctrl_Rep4
Ctrl_Rep2
Ctrl_Rep1
PC2 (18.1%)
PC1 (27.3%)

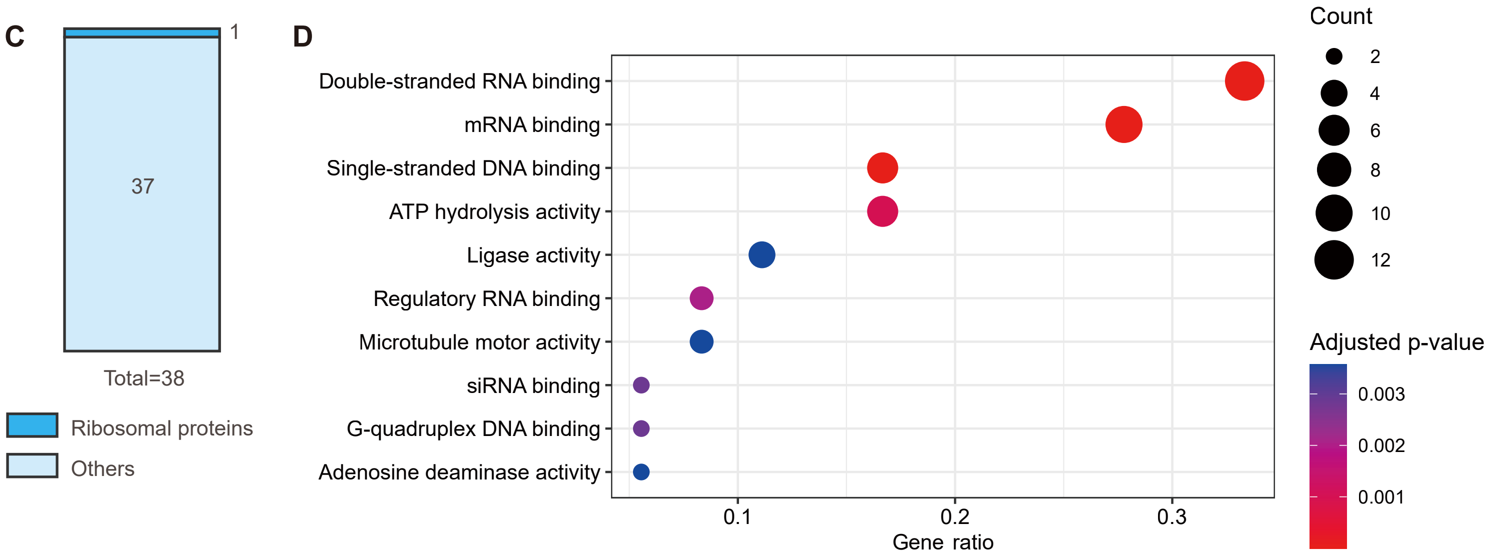


**Figure S3. Mass spectrometry data analysis of poly(I:C) pulled down samples, Related to Figure 3.** (A) Pairwise correlation matrix of proteins detected in each of four biological replicates of poly(I:C) pulled down and control samples by LC-MS/MS. (B) PCA plot showing the distinct characteristics of each experimental group. (C) Number of ribosomal and non-ribosomal proteins in the enriched proteins. (D) A dot plot illustrating the enriched GO terms in molecular function among poly(I:C) interactome, analyzed using the clusterProfiler R package.

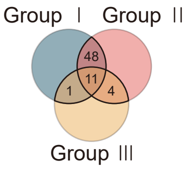


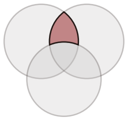

| Gene | Group Ⅰ log$_2$(FC) | Group Ⅱ log$_2$(FC) |
|---|---|---|
| NIFK | 5.749322 | 4.17985 |
| FAM120A | 5.289516 | 2.27006 |
| NAT10 | 3.455037 | 5.30907 |
| RRS1 | 3.121743 | 3.83995 |
| GTPBP1 | 3.101794 | 6.55682 |
| NOP16 | 2.925102 | 3.30531 |
| CCDC86 | 2.909267 | 3.75717 |
| RPF2 | 2.8259 | 5.33858 |
| SRBD1 | 2.812269 | 2.34412 |
| C7orf50 | 2.620101 | 4.29253 |
| YTHDC2 | 2.588107 | 4.50657 |
| PDCD11 | 2.384382 | 7.60389 |
| KRI1 | 2.366952 | 3.66314 |
| C11orf98 | 2.326804 | 4.95941 |
| USP10 | 2.202328 | 2.76475 |
| HNRNPR | 2.179862 | 2.98274 |
| UTP18 | 2.137292 | 2.63391 |
| GNL2 | 2.084256 | 4.17527 |
| ZC3HAV1 | 2.069172 | 3.47621 |
| NOL9 | 1.904099 | 3.2572 |
| GLTSCR2 | 1.842863 | 6.12062 |
| DHX30 | 1.714398 | 4.49709 |
| UPF1 | 1.569153 | 6.93249 |
| MYBBP1A | 1.56017 | 3.27407 |
| NMNAT1 | 1.53313 | 7.20931 |
| EPRS | 1.523144 | 3.5426 |
| GTPBP4 | 1.510063 | 5.66075 |
| LARP1 | 1.509175 | 3.86344 |
| TBL3 | 1.357237 | 3.90217 |
| ZFR | 1.258345 | 2.28191 |
| MOV10 | 1.224216 | 5.99616 |
| EIF6 | 1.212037 | 4.53376 |
| MTDH | 1.206908 | 5.06674 |
| RBM34 | 1.204624 | 7.45403 |
| PTBP1 | 1.202316 | 2.60213 |
| GNL3 | 1.178436 | 3.80406 |
| NPM3 | 1.15409 | 2.62997 |
| RSL1D1 | 1.129534 | 3.02948 |
| KRR1 | 1.117768 | 3.80355 |
| EIF2S2 | 1.112373 | 3.88854 |
| BTF3 | 1.111458 | 3.35558 |
| SRPK1 | 1.101153 | 3.67265 |
| NSA2 | 1.062331 | 3.06405 |
| IGF2BP3 | 1.0618 | 4.55919 |
| NOP2 | 1.060153 | 6.70718 |
| RALY | 1.022763 | 4.01619 |
| CCDC124 | 1.018371 | 3.2902 |
| DDX56 | 1.004732 | 2.86579 |

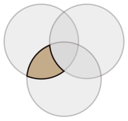

| Gene | Group Ⅰ log$_2$(FC) | Group Ⅲ log$_2$(FC) |
|---|---|---|
| PKR | 2.588877 | 4.946629 |

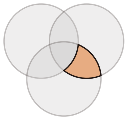

| Gene | Group Ⅱ log$_2$(FC) | Group Ⅲ log$_2$(FC) |
|---|---|---|
| ILF3 | 5.94328 | 1.598356 |
| ILF2 | 2.63903 | 1.403408 |
| TARBP2 | 2.35319 | 2.693842 |
| ADARB1 | 2.05865 | 2.744838 |

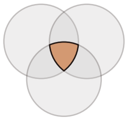

| Gene | Group Ⅰ log$_2$(FC) | Group Ⅱ log$_2$(FC) | Group Ⅲ log$_2$(FC) |
|---|---|---|---|
| STAU2 | 5.297338 | 5.89647 | 4.104683 |
| ZNF346 | 3.355228 | 4.22838 | 8.88317 |
| DHX36 | 3.128029 | 4.38035 | 1.567951 |
| PRKRA | 3.092881 | 6.26501 | 3.874045 |
| L1RE1 | 2.030545 | 3.20107 | 1.461329 |
| ZCCHC3 | 1.787223 | 4.85059 | 1.014778 |
| PURB | 1.675961 | 2.64468 | 7.085686 |
| ADAR | 1.36518 | 8.86358 | 4.511518 |
| STAU1 | 1.132288 | 7.52229 | 2.350547 |
| DHX9 | 1.078274 | 5.87488 | 1.283298 |
| PURA | 1.069166 | 2.91079 | 2.476144 |

**Figure S4. A comparison of the three kinds of interactomes, Related to Figure 3.** Detailed lists of proteins within the overlaps between the K1 interactome, RNase T1-treated K1 interactome, and poly(I:C) interactome, representing the potential dsRNA interactome.

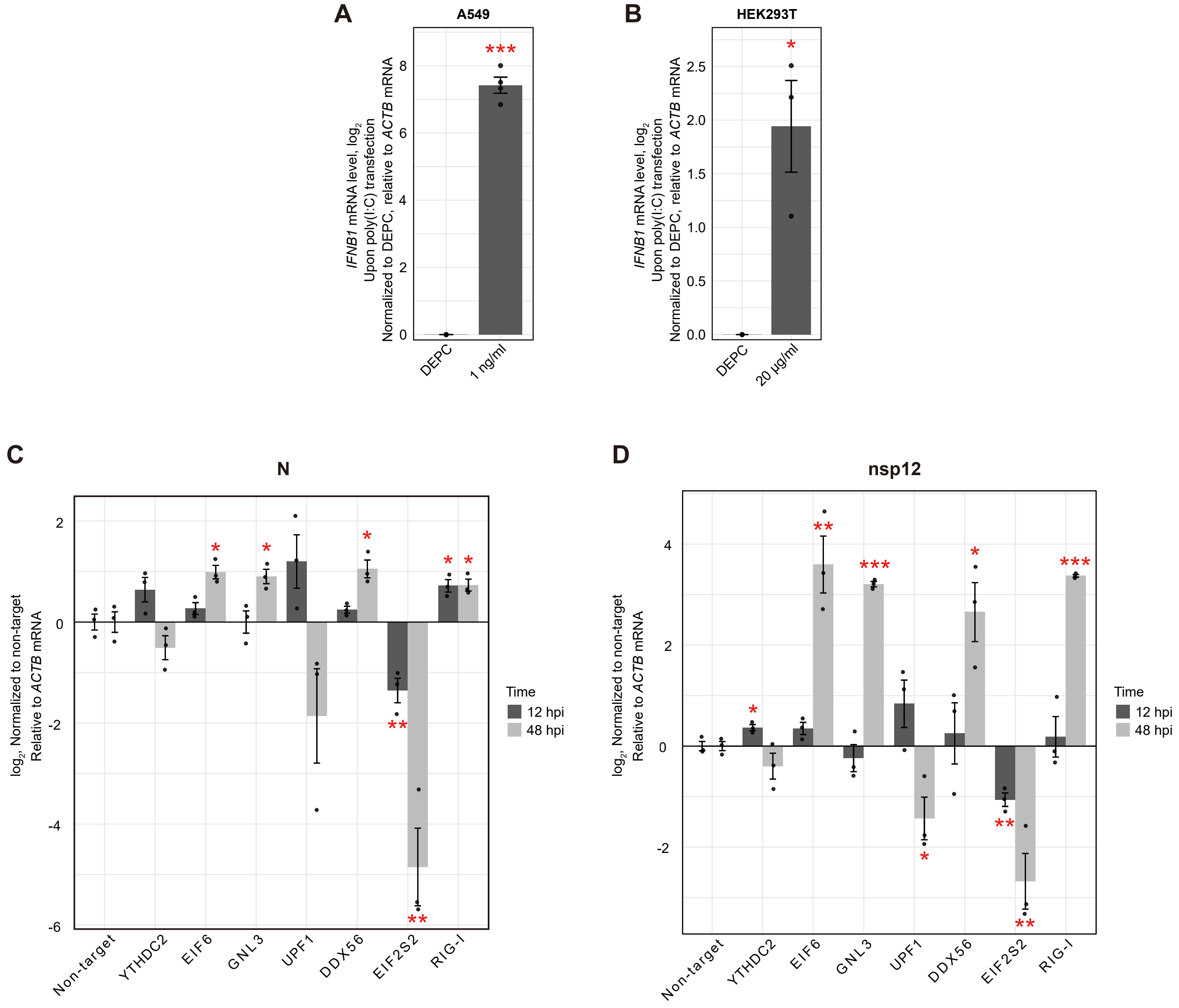


**Figure S5. Characterization of cells transfected with poly(I:C) or OC43 infection, Related to Figure 4.** (A and B) *IFNB1* mRNA induction upon poly(I:C) transfection in A549 (A) and HEK293T (B) cells. An average of three biological replicates is shown with error bars denoting s.e.m. Statistical significances were calculated using two-sided Student's t tests; *p < 0.05 and ***p <0.001. (C and D) N sgRNA (C) and nsp12 gRNA (D) levels measured by RT-qPCR in A549-Cas9 cells depleted with the indicated gene. Viral genes were analyzed at 12 and 48 hpi. An average of three biological replicates are shown with error bars denoting s.e.m. Statistical significances were calculated using two-sided Student's t tests; *p < 0.05, **p < 0.01, and ***p <0.001.
