## Supplementary material for "Cellular dsRNA interactome reveals the regulatory map of exogenous RNA sensing": Key resources table

| REAGENT or RESOURCE | SOURCE | IDENTIFIER |
| --- | --- | --- |
| Antibodies | | |
| double-stranded RNA Monoclonal Antibody (K1) | English and Scientific Consulting Kft | Cat# 10020500; RRID: AB_2756865 |
| Mouse (E5Y6Q) mAb IgG2a Isotype Control | Cell Signaling Technology | Cat# 61656; RRID: AB_2799613 |
| GAPDH (6C5) mouse monoclonal antibody | Santa Cruz Biotechnology | Cat# sc-32233; RRID: AB_627679 |
| GAPDH (D16H11) XP® Rabbit mAb | Cell Signaling Technology | Cat# 5174; RRID: AB_10622025 |
| Anti-Beta Tubulin (D3U1W) Mouse mAb antibody | Cell Signaling Technology | Cat# 86298; RRID: AB_2715541 |
| β-Tubulin (9F3) Rabbit mAb | Cell Signaling Technology | Cat# 2128; RRID: AB_823664 |
| ADAR1 (D7E2M) Rabbit mAb | Cell Signaling Technology | Cat# 14175; RRID: AB_2722520 |
| PKR (D7F7) Rabbit mAb | Cell Signaling Technology | Cat# 12297; RRID: AB_2665515 |
| STAU1 antibody | Abcam | Cat# ab73478; RRID: AB_1641030 |
| UPF1 | Cho et al.^81^ | <https://doi.org/10.1073/pnas.1409612112> |
| Rabbit anti-LARP1 Ab, Affinity Purified | Bethyl Laboratories | Cat# A302-087A; RRID: AB_1604274 |
| Rabbit anti-LARP4 Ab, Affinity Purified | Bethyl Laboratories | Cat# A303-900A; RRID:AB_2620250 |
| DECR1 Polyclonal Antibody | Invitrogen | Cat# PA5-80549; RRID: AB_2787851 |
| ELAVL1 (D9W7E) Rabbit mAb #12582 | Cell Signaling Technology | Cat# 12582; RRID: AB_2797964 |
| Anti-Nucleostemin antibody | Abcam | Cat# ab70346; RRID: AB_1269600 |
| Rig-I (D14G6) Rabbit mAb | Cell Signaling Technology | Cat# 3743; RRID: AB_2269233 |
| Bacterial and virus strains | | |
| HCoV-OC43 | Korea Bank for Pathogenic Viruses | KBPV-VR-8 |
| Chemicals, peptides, and recombinant proteins | | |
| AccuNanoBead™ Protein A Magnetic Nanobeads, size 400nm | Bioneer | Cat# TA-1022-5 |
| NP-40 | Biosolution | Cat# BN015 |
| RNase T1 | Worthington | Cat# LS01490; CAS: 9026-12-4 |
| Protease Inhibitor Cocktail Set III, Animal-Free | Calbiochem | Cat# 535140 |
| BSA | RMBIO | Cat# BSA-BSH |
| Recombinant RNase Inhibitor | Takara | Cat# 2313A |
| Poly(I:C) (HMW) Biotin | InvivoGen | Cat# tlrl-picb |
| Urea | Sigma | Cat# U5378 |
| Seppro™ Ammonium Bicarbonate Buffer | Sigma | Cat# S2454 |
| DL-Dithiothreitol | Sigma | Cat# 43819 |
| Iodoacetamide | Sigma | Cat# I1149 |
| Mass Spectrometry Grade Proteases | Thermo Scientific | Cat# 90057 |
| Calcium chloride (CaCl₂) solution | Sigma | Cat# 21115 |
| Discovery® DSC-18 SPE Tube | Supelco | Cat# 52601-U |
| Acetonitrile (ACN) | Supelco | Cat# 1.00029 |
| Trifluoroacetic acid (TFA) | Sigma | Cat# T6508 |
| Formic Acid (FA), LC-MS Grade, | Thermo Scientific | Cat# 28905 |
| Opti-MEM Reduced Serum Medium | Gibco | Cat# 31985070 |
| Polybrene Infection / Transfection Reagent | Sigma | Cat# TR-1003-G |
| Blasticidin | Gibco | Cat# A1113903 |
| Puromycin | InvivoGen | Cat# ant-pr-1 |
| polyinosinic–polycytidylic acid sodium salt | Sigma | Cat# P1530 |
| TRIzol | Ambion | Cat# 15596026 |
| DMEM, High glucose | Welgene | Cat# LM001-05 |
| RPMI1640 | Welgene | Cat# LM011-01 |
| 0.05% Trypsin-EDTA | Welgene | Cat# LS015-01 |
| Fetal Bovine Serum, qualified, United States | Gibco | Cat# 26140079 |
| GlycoBlue Coprecipitant | Invitrogen | Cat# AM9516 |
| Acid-Phenol:Chloroform, pH 4.5 | Invitrogen | Cat# AM9722 |
| DEPC-DW | Bioneer | Cat# C-9030 |
| Recombinant DNase I (RNase-free) | Takara | Cat# 2270A |
| SensiFAST™ SYBR® Lo-ROX Kit | Bioline | Cat# BIO-94020 |
| RevertAid Reverse Transcriptase | Thermo Scientific | Cat# EP0442 |
| Critical commercial assays | | |
| IFN beta Human ELISA Kit | Thermo Scientific | Cat# 414101 |
| Lipofectamine 3000 Transfection Reagent | Invitrogen | Cat# L3000075 |
| Cell Counting Kit-8 (CCK-8) | Dojindo | Cat# CK04-13 |
| NP-40, for LC-MS/MS | VWR | Cat# 97064-730 |
| KOD -Multi & Epi- | TOYOBO | Cat# KME-101 |
| Expin^TM^ PCR SV | GeneAll | Cat# 103-102 |
| Deposited data | | |
| Raw and analyzed proteomic data | This paper | PRIDE: PXD053100 |
| Experimental models: Cell lines | | |
| Human: HEK293T | ATCC | Cat# CRL-3216; RRID: CVCL_0063 |
| Human: A549 | ATCC | Cat# CCL-185; RRID: CVCL_0023 |
| Human: RD | KCLB | Cat# 10136 |
| Human: HCT-8 | KCLB | Cat# 10244 |
| Oligonucleotides | | |
| Random primers | Thermo Fisher Scientific | Cat# 48190-011 |
| sgRNAs for CRISPR-Cas9 mediated knockout; Table S4 | Bioneer | Table S4 sgRNAs |
| Primers for DNA sequencing; Table S4 | Bionics | Table S4 Primers for DNA sequencing |
| Primers for RT-qPCR; Table S4 | Macrogen | Table S4 Primers for RT-qPCR |
| Recombinant DNA | | |
| pMD2.G | Addgene | Cat# 12259; RRID: Addgene_12259 |
| psPAX2 | Addgene | Cat# 12260; RRID: Addgene_12260 |
| lentiGuide-Puro | Addgene | Cat# 52963; RRID: Addgene_52963 |
| lentiCas9-Blast | Addgene | Cat# 52962; RRID: Addgene_52962 |
| Software and algorithms | | |
| Graphpad Prism 9 | Graphpad software | https://www.graphpad.com/ |
| R (v4.2.1) | R Core Team | https://www.r-project.org/ |
| clusterProfiler package (v4.4.4) | Wu et al.^82^ | <https://doi.org/10.1016/j.xinn.2021.100141> |
| Image Lab | Bio-rad | N/A |
| Illustrator | Adobe | N/A |
